# Molecular Expression Profiles of Morphologically Defined Hippocampal Neuron Types: Empirical Evidence and Relational Inferences

**DOI:** 10.1101/633883

**Authors:** Charise M. White, Christopher L. Rees, Diek W. Wheeler, David J. Hamilton, Giorgio A. Ascoli

## Abstract

Gene and protein expressions are key determinants of cellular function. Neurons are the building blocks of brain circuits, yet the relationship between their molecular identity and the spatial distribution of their dendritic inputs and axonal outputs remain incompletely understood. The open-source knowledge base Hippocampome.org amasses such transcriptomic data from the scientific literature for morphologically defined neuron types in the rodent hippocampal formation: dentate gyrus (DG), CA3, CA2, CA1, subiculum (SUB), and entorhinal cortex (EC). Positive, negative, or mixed expression reports were initially obtained from published articles directly connecting molecular evidence to neurons with known axonal and dendritic patterns across hippocampal layers. Here, we supplement this information by collating, formalizing, and leveraging relational expression inferences (REIs) that link a gene or protein expression or lack thereof to that of another molecule or to an anatomical location. With these additional interpretations, we freely release online a comprehensive human- and machine-readable molecular profile for the more than 100 neuron types in Hippocampome.org. Analysis of these data ascertains the ability to distinguish unequivocally most neuron types in each of the major subdivisions of the hippocampus based on currently known biochemical markers. Moreover, grouping neuron types by expression similarity reveals eight super-families characterized by a few defining molecules.

**Significance Statement:** The molecular composition of cells underlies their structure, activity, and function. Neurons are arguably the most diverse cell types with their characteristic tree-like shapes mediating synaptic communication throughout the brain. Biochemical marker data are available online for hundreds of morphologically identified neuron types in the mammalian hippocampus, including expression of calcium-binding proteins, receptors, and enzymes. Here, we augment this evidence by systematically applying logical rules empirically derived from the published literature (e.g. “presence of molecule X implies lack of molecule Y”). The resulting substantially expanded expression profiles provide nearly unique molecular identities for most known hippocampal neuron types while revealing previously unrecognized genetic similarities across anatomical regions and morphological phenotypes.

## Introduction

Neurons expressing different molecular markers have been found in numerous mammalian brain regions, including the hippocampus (Klausberger, 2008; Baude et al., 2007), barrel cortex (Karagiannis et al., 2009), prefrontal cortex (Caballero et al., 2014), entorhinal cortex (Ferrante et al., 2016), striatum (Garas et al., 2016), retina (Shekhar et al., 2016), and olfactory bulb (Eyre et al., 2009). Because many genes are differentially expressed in different cells, they can serve as chemical identifiers for subsets of cell types. Large-scale efforts examining expression of thousands of genes at the single-cell level are underway (Zeng et al., 2012; Tasic et al., 2016; Poulin et al., 2016; Macosko et al., 2015; Zeisel et al., 2015; Chen et al., 2017; Hu et al., 2017; Raj et al., 2018). The big data from these studies will advance significantly our knowledge of the molecular markers controlling neuronal function. However, full elucidation of how the brain works will likely require additional dimensions of information that may not be directly derivable from the transcriptome alone. In particular, synaptic connectivity, which depends at least in part upon the spatial overlap of the axons and dendrites of two neurons, may also be necessary (see, however, Paul et al., 2017).

Some neurons with known axonal-dendritic patterns are associated with particular molecular markers. For example, hippocampal CA1 Oriens/Lacunosum-Moleculare (O-LM) and neocortical Martinotti neurons express the neuropeptide somatostatin (SOM); however, these are not the only SOM-positive cells in their respective brain regions (Leao et al., 2012; McGarry et al., 2010; Markram et al., 2004). Thus, SOM is a biochemical marker for these neurons, but it does not uniquely identify them. To date, no single marker has been found to uniquely identify a morphologically- or electrophysiologically-distinct neuron type. Rather, uniquely identifying neuron types requires molecular expression information for several markers.

The potential implications of the ongoing transcriptomics revolution are unambiguously recognized in the neuroscience community and the advent of high-throughput single cell sequencing techniques such as DropSeq (Macosko et al., 2015) is rapidly delivering on the promise of much needed large-scale datasets. However, scientific consensus has yet to be reached as to which genes should be considered as relatively stable and informative with respect to neuron types and which may be more dependent on the organism’s state or individual life history. In particular, two critical questions remain open: based on current knowledge of molecular expression, how many morphologically or functionally distinguished neuron types can we uniquely identify in a given brain region based on biochemical markers? And what is the minimal subset of robust neurochemical markers required for such identification?

Molecular expression information for neurons in the rodent hippocampal formation, one of the most studied mammalian brain systems, is available at Hippocampome.org, a free internet knowledge base that also provides morphological, electrophysiological, and synaptic information (Wheeler et al., 2015). Hippocampome.org groups neurons into types based on their primary neurotransmitter and the presence or absence of their axons and dendrites in the layers of each hippocampal subregion.

The present work focuses on the molecular marker profiles for the 122 neuron types in Hippocampome.org version (v.) 1.3. We first profiled each type with biochemical expression data explicitly linked to the identified morphology in the published literature. Then we augment this neuron type-specific evidence by leveraging relational expression inferences (REIs), which link the presence or absence of a gene or protein to that of another molecule (e.g. co-expression or mutual exclusion), or the particular anatomical location of the soma in the hippocampus. Lastly, we explore the ability of these molecular profiles to distinguish neuron types, as well as to suggest molecular families of neuron types. Importantly, the public online release of these augmented profiles in both human- and machine-readable formats enables many additional analyses that may help elucidate the function of both neuron types and genes.

## Materials and Methods

### Obtaining direct molecular expression evidence

Over 1,000 peer-reviewed publications were data mined for molecular expression information in neurons of the hippocampal formation of rats or mice. From this literature, 237 papers showed data indicating expression or lack of expression via immunohistochemistry, *in situ* hybridization, or single cell RT-PCR in neurons matching the descriptions of Hippocampome.org types based on neurite location and putative neurotransmitter. In addition, we used data from the Allen Gene Expression Atlas as the basis for assigning the presence or absence of certain molecular markers in the principal neuron types DG Granule, CA1 Pyramidal, CA2 Pyramidal, and CA3 Pyramidal (Hamilton et al., 2017). All of these data are considered neuron type-specific evidence and lead to interpretations of positive or negative expression in Hippocampome.org. In certain cases, contrasting evidence exists for a given marker being present and absent in a neuron type (“mixed” expression). This could point to the existence of neuronal subtypes or to differences between species (e.g. mouse vs. rat), protocol (e.g. gene vs protein detection), subcellular localization (e.g. somatic vs neuritic) or unknown factors. Each piece of evidence was annotated with essential metadata and all relevant excerpts (text, figures, and references) were stored in and made available through Hippocampome.org (hippocampome.org/markers).

### Obtaining relational expression inferences (REIs)

REIs were data mined from the 237 articles that provided neuron type-specific interpretations. From these, 742 distinct inferences were obtained under three broad categories. In the first, if expression data indicates that an entire hippocampal formation layer or subregion is negative for a given marker, Hippocampome.org neuron types with soma locations exclusively in the encompassed parcel are inferred to be negative for that marker. The remaining categories of REIs capitalize on expression relationships between two markers. One occurs when the literature indicates that expression of a given marker implies co-expression of another marker. This co-expression relation, annotated in Hippocampome.Org as “probability-positive,” is directional, because the presence of the second marker does not always imply that the first one is expressed as well. When data demonstrated actual bidirectional co-expression, we created two probability-positive REIs. The last category consists of a literature indication that the presence of a given marker implies the absence of another marker. This “probability-negative” exclusivity relation is also generally directional: as with the probability-positive REIs, the cases in which presence of the second marker also entails absence of the first (mutual exclusion) are represented by two REIs.

These latter two REI categories are formal IF/THEN statements; consequently, each has a logical contrapositive. In the contrapositives, the position of the markers in the IF/THEN statements is swapped, and the expression interpretation for each is reversed (i.e. positive becomes negative and vice versa). For co-expression REIs, which are of the form “if marker 1 is positive, then marker 2 is positive,” the contrapositive states “if marker 2 is negative, then marker 1 must be negative”. Similarly, for exclusion REIs of the form “if marker 1 is positive, then marker 2 is negative,” the contrapositive is “if marker 2 is positive, then marker 1 must be negative.”

### Evaluating usability of REIs

The REIs were collected from articles that presented the data in different forms, ranging from those that provided frequency of the event and sample size to those that consisted of a text statement describing an expression relationship. We established rules for using each of these to avoid including and propagating inferences based on weak evidence. In formulating the rules, we took into account our confidence in the data and the resulting number of usable REIs. To clarify, if we only accepted REIs that were true in 500 neurons 100% of the time, very limited knowledge could be gained. Conversely, if we accepted those that sampled 3 or more cells and were true >50% of the time, we would have many inferences but diminished confidence. The rules that we established balance these factors.

The layer negative (category one) REIs were considered usable if the layer was completely negative for a molecular marker. Any evidence of neuronal expression of the given marker in that layer disqualified the REI.

For probability-positive and probability-negative inferences, the strength of the REI depends on the sample size (n) and the frequency for which the expression relationship holds (% times true: PTT). When both of these were reported, we calculated the 95% binomial confidence interval using the Clopper-Pearson method (Clopper & Pearson, 1934). We set the threshold for usability at a Clopper-Pearson interval lower bound of 67%, indicating 95% confidence that the REI is true for at least two-thirds of the neurons in the population covered by the REI. The actual average PTT for the REIs included using this criterion data was 93.5% (range 72.0 - 100%), and more than 70% of these REIs had PTT ≥90%.

For many inferences, either the sample size was not provided or the PTT was very high but the sample size was low. We established the following provision for using this material. Inferences with a PTT of 90% or greater were included regardless of sample size. Some evidence for an REI was in the form of text statements that provided no n’s and only implying a PTT (e.g. “No SS [somatostatin] immunoreactivity was found in any 5-HT3R-expressing neurons, although single SS-immunopositive or 5-HT3R-expressing neurons were identified readily in cortex and hippocampus” (Morales & Bloom, 1997)). In these cases, the REIs were used when the statements implied a PTT of 100%.

### Experimental design

We developed MATLAB code for applying the usable REIs to the Hippocampome.org knowledge base. In order to leverage information chaining, the algorithm applies usable REIs in several iterations, each building off of both direct neuron type-specific evidence and REI-based interpretations obtained in previous applications of the rules. The initial data for REI application are the set of solely positive or negative (i.e. not mixed) neuron type-specific molecular marker interpretations in Hippocampome.org. Subsequent passes applied the REIs to REI-based interpretations obtained in earlier iterations if not resulting in a suggestion of mixed expression. Markers that became mixed in the iterative applications were discontinued from further REI application. Iteration stopped when application of the set of REIs yielded no new information.

### Statistical analysis and correlation-produced REIs

To uncover possible new relationships between markers in our knowledge base, a 2-by-2 contingency table for each pair of markers was assessed using Barnard’s exact test. This statistical test is the most robust for contingency tables where the row and column totals can vary (Lydersen, Fagerland, & Laake, 2009).

Correlations that were statistically significant (p ≤ 0.05) and were not already present in the usable REIs were added to the REI list. Subsequently, these correlation-produced inferences were applied once to non-mixed expression interpretations based on neuron type-specific and “regular” REIs; however, no additional iterations were carried out using the interpretations derived from the correlation-produced REIs because of the weaker nature of this evidence compared to regular REIs. Specifically, correlation-produced REIs are based on neuron types, and we do not have counts of how many neurons might be in these neuron types or if these types are the only neurons in a subregion. In addition, unlike regular REIs, they are linked to subregions and not hippocampal formation layers.

### Molecular marker usefulness rating for discriminating neuron types

Following the application of REIs, the molecular markers were assessed in terms of their potential usefulness for identifying neuron types. The assessment took into account the number of neuron types with positive or negative expression interpretations and measured up each marker against a hypothetical, optimally informative marker that would be known for all neuron types and split them evenly into positive or negative (i.e. 50% would be positive and 50% negative). The usefulness ratio was calculated by dividing the product of the numbers of positive and negative neuron types for the marker in question by this same product for the optimal marker, which for the 122 neuron types in Hippocampome.org is 3,721 (61*61).

### Cluster analysis to group neuron types by molecular similarity

SPSS TwoStep Cluster analyses were performed on the molecular marker expression data. Both automated and manual modes were tested: in manual mode the user designates the markers used and the number of clusters in the output, whereas in automated mode the program selects the optimal number of clusters and markers to use. These cluster analysis were always unsupervised, meaning that we never ourselves assigned neuron types to specific clusters.

We obtained results for from five to 20 clusters, using from eight to 20 markers or all markers (224 total outputs). The 20 markers used followed the order of their usefulness ratings, namely PV, CR, SOM, mGluR1a, GABA-Aa1, CB, NG, VIP, CCK, CB1, NPY, a-act2, CoupTF II, AR-beta1, 5HT-3, RLN, nNOS, vGluT3, sub P rec, and ErbB4 (see Figure 1 for abbreviations). To assess the quality of the clustering outputs, we compared the between-neuron marker profile similarity within and across clusters. In particular, the ideal output would group neuron types into tight and well-separated clusters. A tight cluster consists of a set of neuron types with very similar marker profiles, whereas two neuron types from well-separated clusters would have very different marker profiles. We quantified the marker profile similarity for a given pair of neuron types as the difference between the number of markers with the same expression (positive in both types or negative in both types) and the number of markers with opposite expression (positive in one type and negative in the other). We then measured the quality of clustering output using the Calinski-Harabaz index, defined as the ratio between the average pairwise similarity within clusters and the average pairwise similarity across clusters (Calinski & Haraasz, 1974). The highest within-cluster/across-cluster average similarity ratio indicated optimal clustering result.

**Figure 1.**
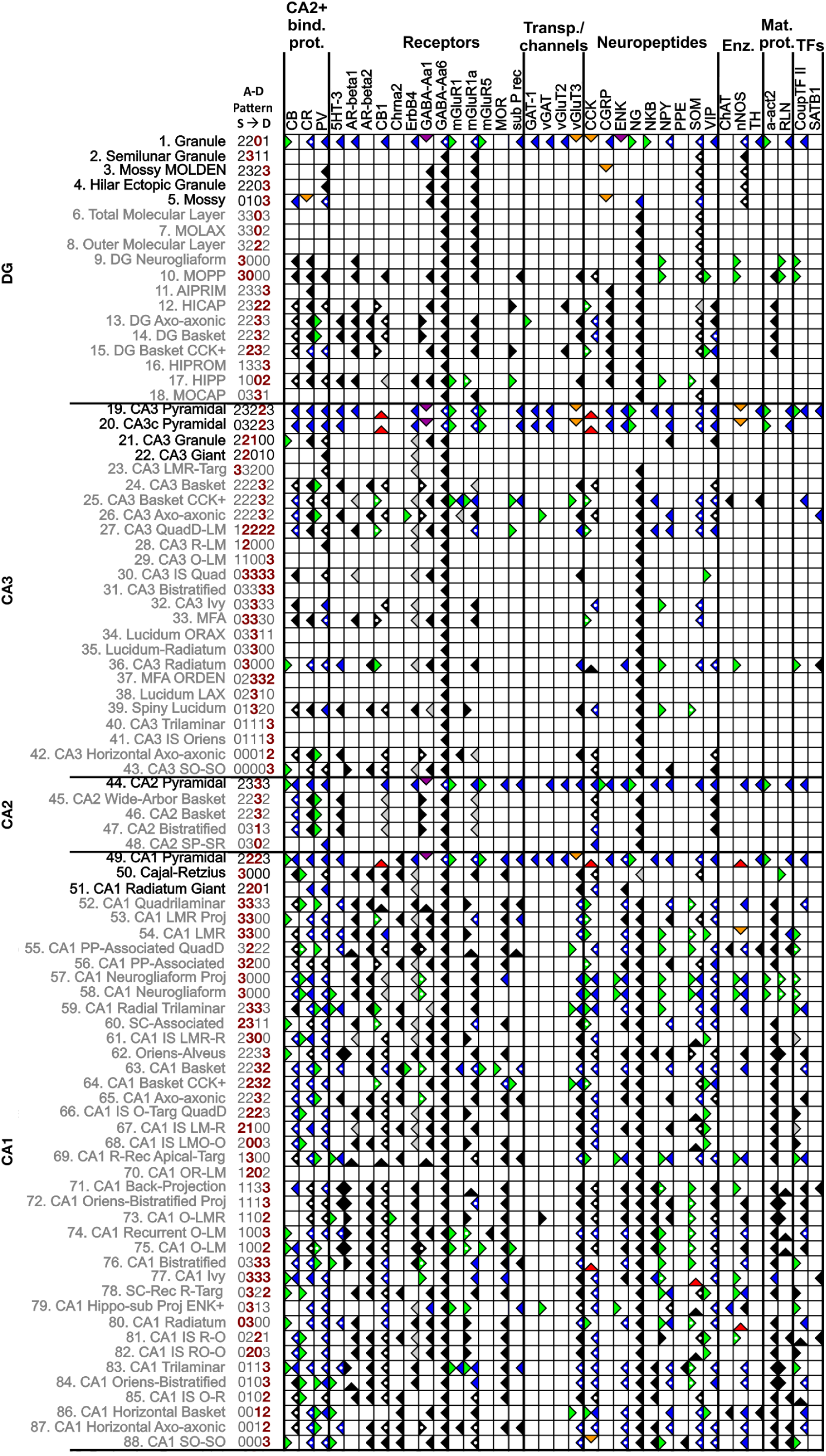
Molecular marker and neuron types in Hippocampome.org v.1.3. This matrix illustrates a subset of the expression information available online (for symbol key, see Table 2). The 36 molecules that have REI-based interpretations and the 88 neuron types in DG, CA3, CA2, and CA1 with two or more REI interpretations are included (putative excitatory and inhibitory neuron types in black and grey fonts, respectively). The defined pattern of axons and dendrites across layers (superficial to deep) for each neuron type is coded numerically beside the name: 1, only axons present; 2, only dendrites present; 3, both axons and dendrites present; 0, neither axons nor dendrites present; red numbers indicate the somatic layers for that neuron type. *<u>Markers:</u>* 5HT-3 (serotonin receptor 3), a-act2 (alpha actinin 2), AR-beta1/2 (andrenergic beta 1/2 receptor), CB (calbindin), CB1 (cannabinoid receptor type 1), CCK (cholecystokinin), Chrna2 (acetylcholine receptor alpha 2 subunit), CGRP (calcitonin gene related peptide), ChAT (choline acetyltransferase), CoupTF II (chicken ovalbumin upstream promoter transcription factor II), CR (calretinin), ENK (enkephalin), GABA-Aa1/6 (gamma aminobutyric acid A alpha1/6 subunit), mGluR1/1a/5 (metabotropic glutamate receptor 1/1a/5), MOR (mu-opioid receptor), NG (neurogranin), NKB (neurokinin B), nNOS (neuronal nitric oxide synthase), NPY (neuropeptide tyrosine), PPE (preproenkephalin), PV (parvalbumin), RLN (reelin), sub P rec (substance P receptor), SOM (somatostatin), TH (tyrosine hydroxylase), vGluT3/2 (vesicular glutamate transporter 2), VIP (vasoactive intestinal polypeptide). Other abbreviations: Enz (enzymes), IS (interneuron specific), Mat (matrix, as in extra- and intracellular matrix), MFA (mossy fiber-associated), PP (perforant path), Proj (projecting), Rec (receiving), SC (Schaffer collateral), Targ (Targeting), TFs (transcription factors).

### Data Availability

A complete list of the “regular” REIs and correlation-produced REIs can be found in the Help menu of Hippocampome.org (hippocampome.org/phpdev/data/REIs.xlsx). In addition, the evidence supporting each expression interpretation can be obtained by clicking the symbols in the molecular marker matrix (hippocampome.org/phpdev/markers.php).

## Results

### Collating molecular marker evidence: neuron type-specific evidence and REIs

The molecular marker interpretations in Hippocampome.org v.1.3 provides information for 97 genes/proteins for which expression or lack of expression was shown in at least one of the 122 morphologically-identified neuron types in the knowledge base (hippocampome.org/markers). This massive endeavor yielded expression information for 894 of the 11,834 entries in the full neuron-type × marker matrix (7.5% filled), as illustrated in Figure 1 for a subset of the available data. These data are represented in the matrix by the (right-facing) green flag (positive expression), (left-facing) blue flag (negative expression) or by a variety of flags indicating mixed expression for different reasons: green and blue (subtypes), orange (species and/or protocol differences), purple (subcellular expression differences, for example soma vs. axons), and red flags (contrasting experimental evidence for unresolved reasons).

Approximately 50 articles describing neuron type-specific molecular evidence in Hippocampome.org also contained REI information, and from them 742 REIs were obtained. Of these, 504 met the usability criteria. The full catalog of REIs includes the contrapositives of the probability positive and probability negative REIs, yielding a total of 988 usable REIs. Approximately half, 477, were applicable to Hippocampome v.1.3 neuron types (Table 1).

**Table 1.**
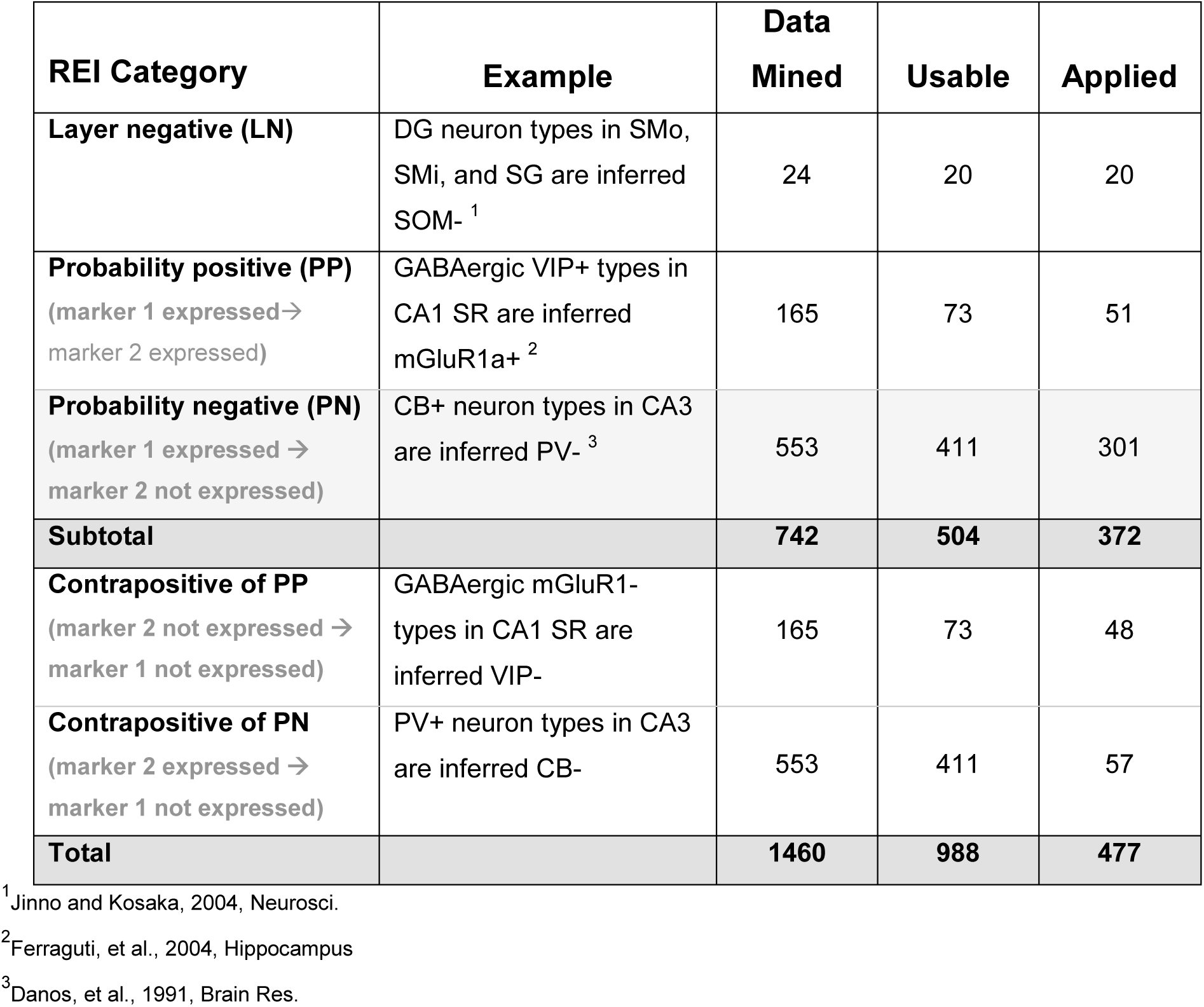
Categories and numbers of REIs

| REI Category | Example | Data Mined | Usable | Applied |
| --- | --- | --- | --- | --- |
| **Layer negative (LN)** | DG neuron types in SMO, SMi, and SG are inferred SOM-<sup>1</sup> | 24 | 20 | 20 |
| **Probability positive (PP)**<br>(marker 1 expressed → marker 2 expressed) | GABAergic VIP+ types in CA1 SR are inferred mGluR1a+<sup>2</sup> | 165 | 73 | 51 |
| **Probability negative (PN)**<br>(marker 1 expressed → marker 2 not expressed) | CB+ neuron types in CA3 are inferred PV-<sup>3</sup> | 553 | 411 | 301 |
| **Subtotal** |  | **742** | **504** | **372** |
| **Contrapositive of PP**<br>(marker 2 not expressed → marker 1 not expressed) | GABAergic mGluR1- types in CA1 SR are inferred VIP- | 165 | 73 | 48 |
| **Contrapositive of PN**<br>(marker 2 expressed → marker 1 not expressed) | PV+ neuron types in CA3 are inferred CB- | 553 | 411 | 57 |
| **Total** |  | **1460** | **988** | **477** |

<sup>1</sup>Jinno and Kosaka, 2004, Neurosci.
<sup>2</sup>Ferraguti, et al., 2004, Hippocampus
<sup>3</sup>Danos, et al., 1991, Brain Res.

### REIs add significantly to molecular expression data for neuron types

In the first of the iterative applications of the REIs, all of the usable inferences were assessed against the neuron type-specific evidence indicating positive or negative, but not mixed, expression (green flags and blue flags in Figure 1). When the criterion for an REI was satisfied, the REI was applied. If the REI provided new information, a black flag (right-facing, positive; left-facing, negative) was placed in the matrix. If it supported an already present interpretation, a white dot was placed in the existing flag, representing an increased level of confidence in that interpretation (Figure 2).

**Figure 2.**
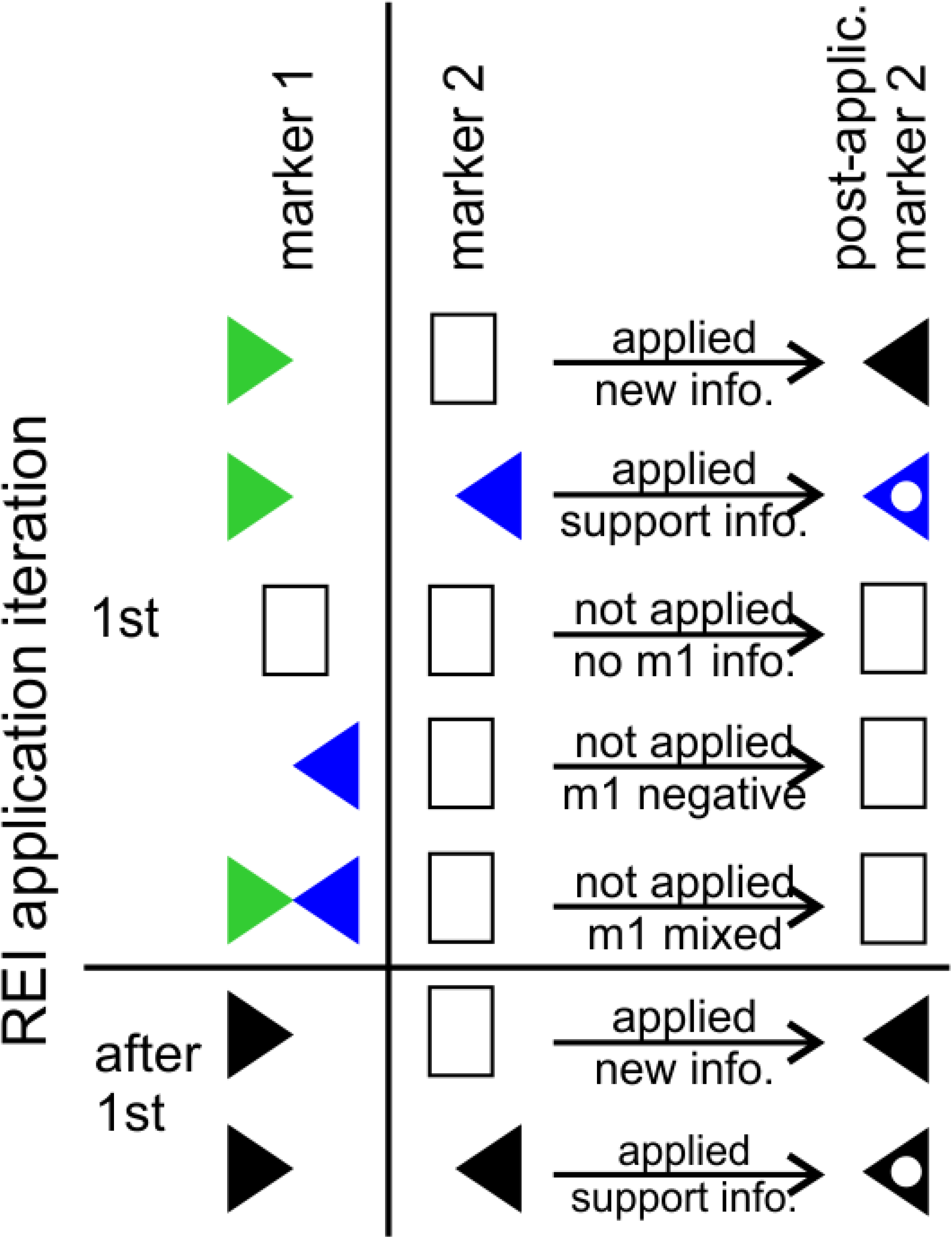
Iterative application protocol for probability REIs. In the first iteration, all usable REIs were assessed against the neuron type-specific information. In subsequent iterations, they were assessed against REI-based interpretations. In this example, a probability negative REI (i.e. if marker m1 is positive, m2 is inferred negative) is assessed against all of the m1 data. The REI is applicable to any neuron type that is positive for m1, but is not applicable to negative or mixed interpretations. If m1 is positive and m2 is unknown, a black flag facing left will be placed in m2, indicating inferred negative expression. If m2 already has an interpretation and the REI supports it, a white dot is placed in the existing flag. If the REI interpretation conflicts with an existing interpretation, no flag is added. In the next iteration, any new black flags are assessed. The iterations continue until no more information is added to the molecular marker matrix.

In subsequent iterations, the assessment was performed on REI-based interpretations and new black flags or white dots in existing black flags were added to the matrix. If contradictory information arose in the same iteration, both the positive and negative information was added, creating a mixed interpretation. In these cases, we evaluated the evidence to determine whether the conflict suggested subtypes, could be attributed to species/protocol differences, or could not be resolved (Table 2). All of the 21 mixed occurrences in these analyses belonged to the latter category.

**Table 2.**
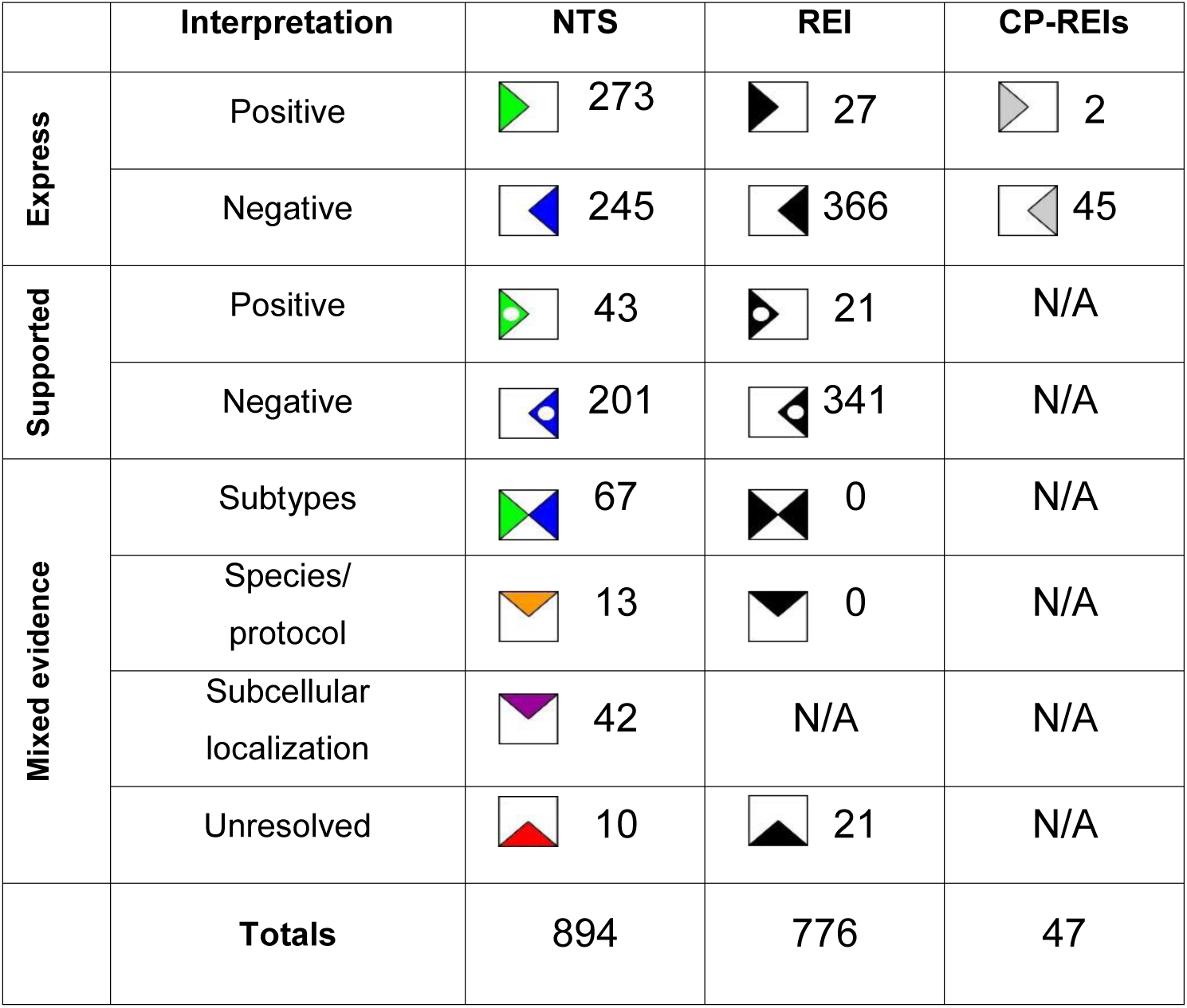
Molecular marker expression interpretations resulting from neuron type-specific evidence (NTS), REIs, and correlation-produced REIs.

|  | Interpretation | NTS | REI | CP-REIs |
| --- | --- | --- | --- | --- |
| **Express** | Positive | 273 | 27 | 2 |
|  | Negative | 245 | 366 | 45 |
| **Supported** | Positive | 43 | 21 | N/A |
|  | Negative | 201 | 341 | N/A |
| **Mixed evidence** | Subtypes | 67 | 0 | N/A |
|  | Species/<br>protocol | 13 | 0 | N/A |
|  | Subcellular<br>localization | 42 | N/A | N/A |
|  | Unresolved | 10 | 21 | N/A |
|  | **Totals** | 894 | 776 | 47 |

The 20 applicable layer-negative REIs provided information for 7 molecular markers, and the probability positive/negative and their contrapositive REIs provided information for an additional 29 (Figures 1 and 3). Together, they increase expression knowledge for molecular markers falling into the categories of calcium-binding proteins, receptors, transporters and channels, neuropeptides, enzymes, matrix proteins, and transcription factors in neuron types across all hippocampal formation subregions.

**Figure 3.**
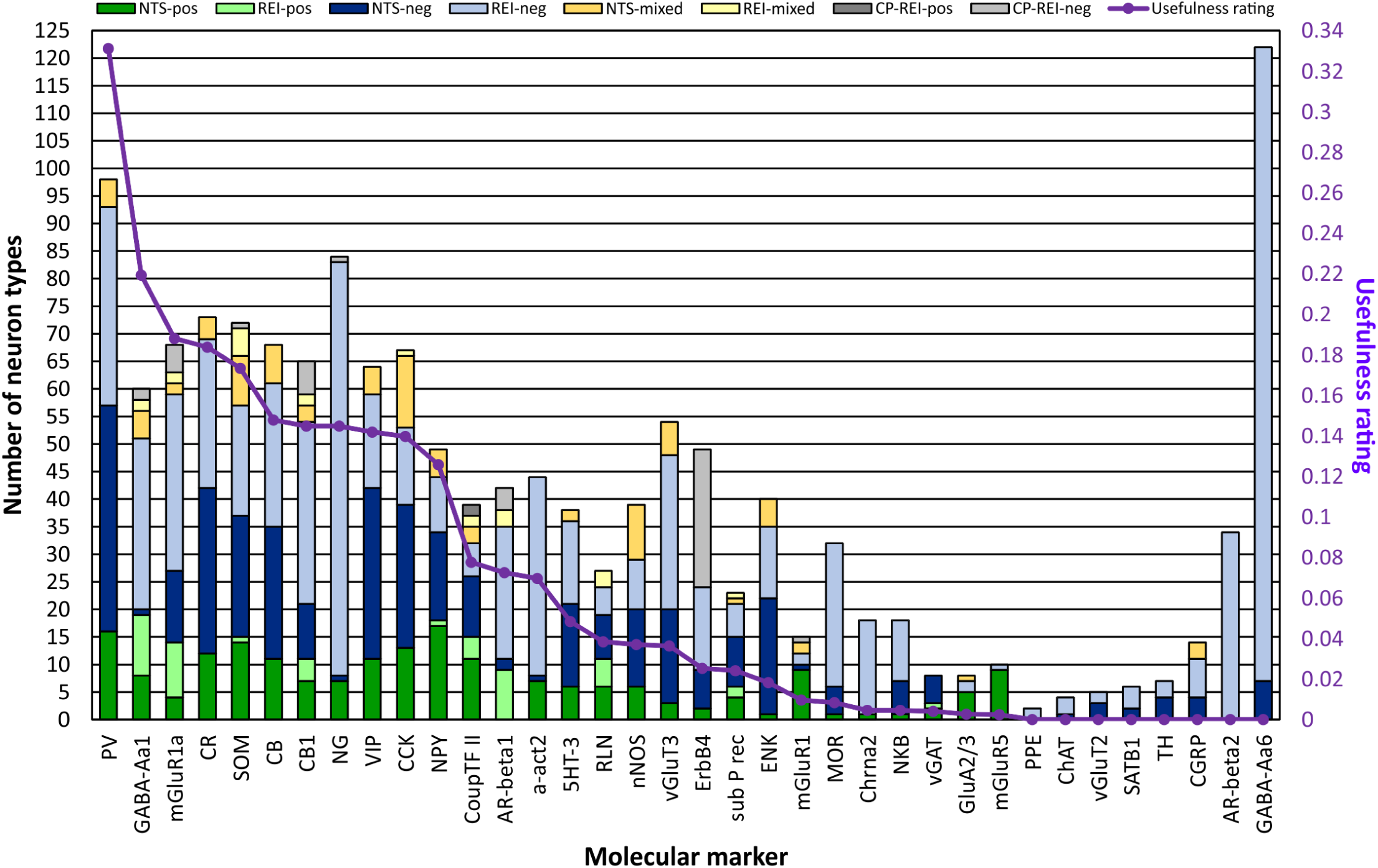
REIs increase expression information for 36 markers and enhance the usefulness in distinguishing neuron types. For 15 of these 36 markers, the REI- and correlation-produced (CP) REI-based interpretations more than double the number of neuron types with interpretations from neuron types-specific (NTS) evidence alone. Moreover, REIs are the only source of interpretations for some markers. In evaluating the usefulness of each marker in distinguishing rodent hippocampal neuron types, we considered the number of positive- and negative-expressing neuron types in comparison to a hypothetical optimal marker. Although the number of neuron types that are negative for PV expression exceeds the number that are positive, PV is first in usefulness because its expression is known for the most neuron types.

The molecular marker with information for the most neuron types is GABA-Aa6 (Figure 3). Of the 36 markers presented, it is unique because data show that it is not expressed in the hippocampal formation (Hortnagl et al., 2013); consequently, all of the neuron types are negative for it. The same interpretations would be made for all genes not transcribed in the hippocampal formation. Although knowledge of these universally negatively expressed genes potentially is insightful for elucidating the behavior of neuron types, they are not beneficial in distinguishing neuron types. Consequently, we are not presenting more of them in this work. Excluding GABA-Aa6, the other 35 markers with REI-based interpretations added information to 102 neuron types, of which the 88 located in DG, CA3, CA2 or CA1 are shown in Figure 1.

### Using the compendium of molecular marker information to identify new REIs

After filling the marker matrix with neuron type-specific and REI-based marker expression interpretations data, we performed correlation analyses that compared the expression of each molecular marker to all of the other markers to identify potentially new REIs not discovered through literature mining. Thirteen correlations reached significance (p ≤ 0.05). These included ones suggesting that the neuropeptide NG and the extracellular matrix protein RLN have mutually exclusive expression, and that expression of ErbB4 co-localizes with PV (full list at hippocampome.org/phpdev/data/REIs.xlsx). Application of the correlation-produced REIs resulted in two new positive and 45 new negative interpretations (Table 2 and grey flags in Figure 1). The correlation-produced REIs add the most information to ErbB4, but also contribute to GABA-Aa1, mGluR1a, SOM, CB1, CoupTF II, and AR-beta 1 (Figure 3). Altogether, leveraging various types of REIs almost doubled the amount of available molecular marker information for hippocampal neuron types, generating 823 new expression interpretations for a grand-total of 1717 flags or >14.5% of the full molecular marker matrix of 122 neuron types by 97 markers (Figure 1, Table 2, and hippocampome.org/phpdev/markers.php). Moreover, REIs provide supporting data for 244 neuron type-specific and 362 REI-based interpretations.

### Metric to evaluate the usefulness of individual markers in distinguishing neuron types

One potential use of molecular markers is to distinguish neuron types. Among the 36 molecules listed in Figures 1 and 3 are some of the most investigated proteins in hippocampal research; however, their usefulness in identifying neurons varies depending on their ratios of positive to negative interpretations and on the number of neuron types with positive or negative expression. A hypothetical optimal marker would have information for all neuron types and have a positive/negative ratio of one. The data from each of the 36 markers when compared to this optimal marker fall substantially below the ideal (Figure 3). Current data suggest that PV is the most useful in distinguishing rodent hippocampal neuron types with a usefulness rating of ∼0.33, followed by GABA-Aa1 (0.22), mGluR1a (0.19), CR (0.18), and SOM (0.17).

The trend for all of the data is that the number of neuron types that are negative for a particular marker (light or dark blue or light grey bars in Figure 3) is greater than those that are positive. CoupTF II and RLN have more balanced positive/negative ratios, 1.0 and 0.85, respectively; however, they have lower usefulness ratings (0.08 and 0.04, respectively) because their expression is known for fewer neuron types. If they maintain the same balanced ratios when data for more neuron types are published, these markers may become the most useful in distinguishing rodent hippocampal neuron types.

### Using molecular marker information to distinguish neuron types

#### Pairwise comparisons

With the amassed expression interpretations, each of the Hippocampome.org neuron types has a molecular marker expression profile. We analyzed these profiles to assess their uniqueness among the neuron types. Specifically, the profile of each neuron type was compared to those of all of the other neuron types (Figure 4A). The results of these pairwise comparisons are shown in Figure 4B. Using only their molecular marker profiles, there is insufficient information to conclude that the molecular profile of any neuron type is unique among the 122 types because some neuron types have expression interpretations for a single or very few markers; specifically, only 40.4% of the comparisons show different profiles (squares not grey or white in Figure 4B).

**Figure 4.**
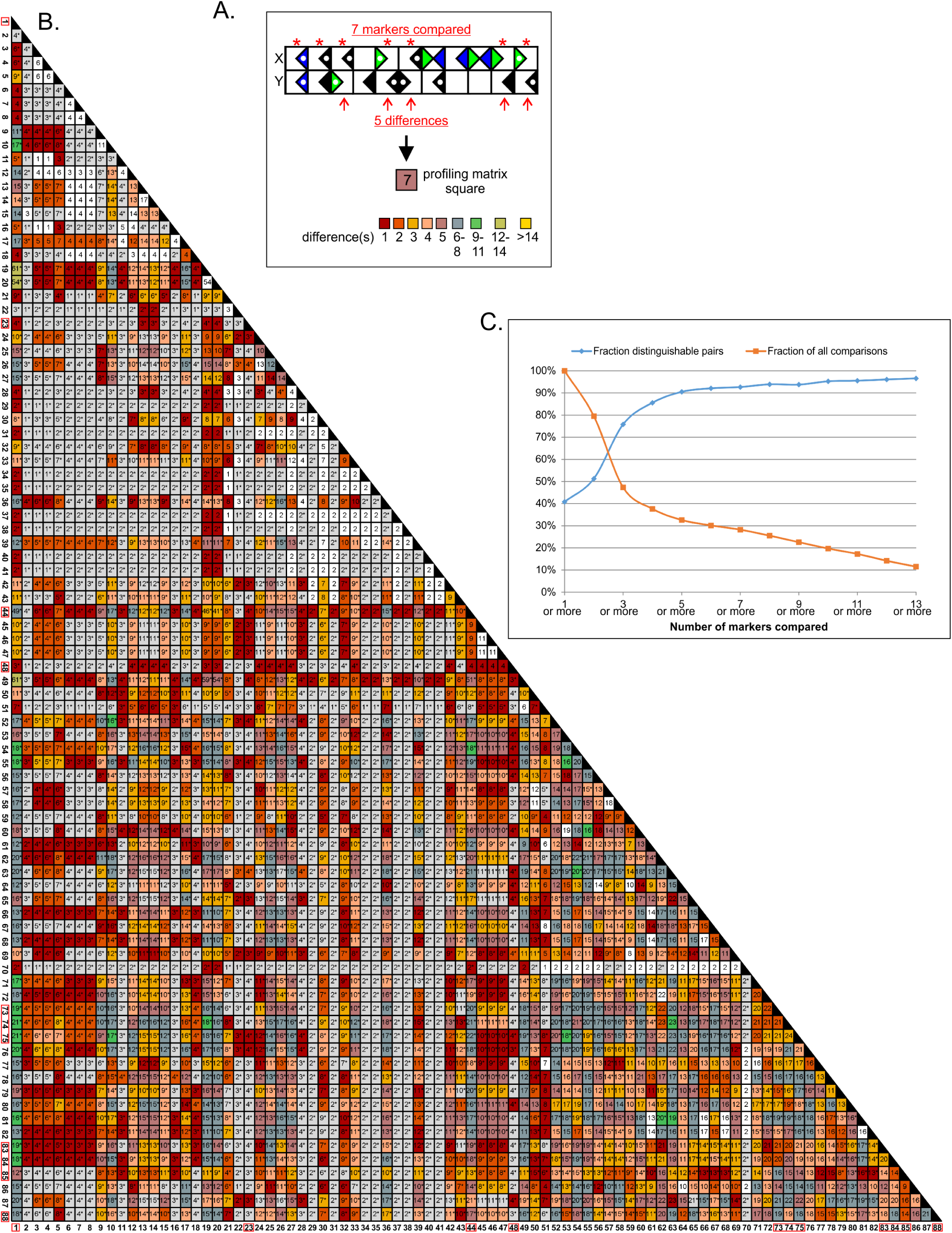
Finding unique molecular marker profiles in the 122 neuron types. **A.** Each pairwise comparison includes any marker that has evidence for both neuron types considered indicating either positive or negative, but not mixed, expression. The number of included markers is placed in the appropriate square of the profiling matrix (panel B) while the count of differently expressed markers is indicated by the color of the square. **B.** Profiling matrix for the 88 neuron types shown in Fig. 1, but the entire matrix has 122 types (yielding 7,381 possible pairwise comparisons). Of the 5,885 pairs with at least one marker in common in addition to universally negative GABA-Aa6 (squares with ≥2), 50% show different profiles (non-white/grey squares). When soma location is factored in, the proportion of distinguishable pairs rises to 94.4%, and 11 neuron types have unique profiles (red boxes around type numbers). **C.** The number of distinguishable neuron type pairs increases with the number of markers with known expression for both types: when 8 or more markers can be compared, ≥95% of pairs can be distinguished; however, less than 30% of the pairs meet or exceed this threshold.

In the strictest terms, however, each neuron type can be distinguished from most of the others on the basis of the location(s) of its soma (indicated by grey squares and asterisks in the non-grey squares in Figure 4B). To illustrate, a researcher looking in an intact tissue slice for CA1 O-LM neurons, which have their dendrites in stratum oriens and the bulk of their axons in stratum lacunosum-moleculare (Hippocampome.org neurite pattern 1002), would likely not confuse them with DG HIPP, which have the same neurite pattern but are located in a different subregion. However, this could be problematic when dissociated neurons are investigated. Taking into account soma location along with marker profiles, 94.4% of the neuron type pairs can be differentiated (non-white squares in Figure 4B). Based on the combined information of soma location and marker profiles, 11 neuron types can be distinguished from all other types (red boxes around neuron type numbers in Figure 4B). Interestingly, CA1 O-LM 1002 and DG HIPP 1002 neuron types can be distinguished not only by having somata in different subregions, but also by molecular profiles with data suggesting that they may differ in the expression of three markers (Figure 1), namely PV (Kosaka et al., 1987; Katona, Acsady, & Freund, 1999; Tukker et al., 2007), GABA-Aa1 (Gao & Fritschy, 1994; Viney et al., 2013), and AR-beta1 (Cox, Racca, & LeBeau, 2008).

In the pairwise comparisons, only non-mixed interpretations were used. The overall average number of molecular markers considered in each analysis (Figure 4B, numbers in squares) was five (range 1-56) excluding identity comparisons (i.e. a neuron type compared to itself). This average rises to nine when only comparisons that returned distinguishable types are assessed. As the number of molecular markers compared increases, so does the percentage of distinguishable pairs (Figure 4C). When eight or more markers are compared, ≥95% of the pairwise comparisons return distinguishable neuron types; however, <30% of the comparisons reach that mark.

#### Identifying neuron types in a hippocampal subregion and layer

To test the power of the Hippocampome.org molecular marker dataset, we attempted to uniquely identify all of the neuron types in CA1 stratum oriens (SO) based on the Hippocampome.org collection of molecular markers. We chose a layer in CA1 because it has the most information both in terms of number of neuron types and marker information. In Hippocampome.org v.1.3, 17 neuron types have somata in CA1 SO (Figure 5).

**Figure 5.**
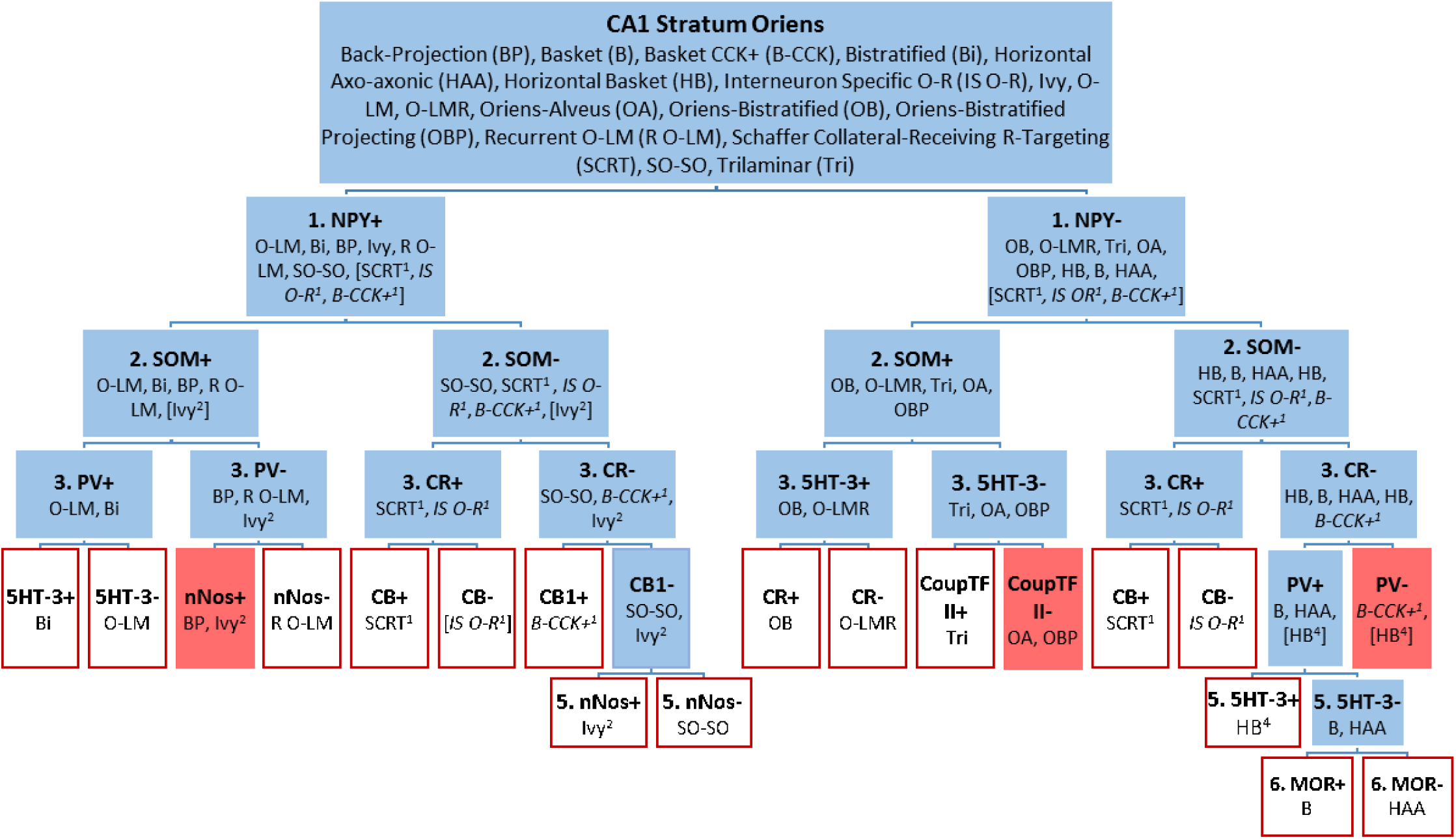
Most of the 17 neuron types with soma in CA1 SO have unique molecular marker profiles. Neuron types are divided into positive or negative expression for each marker queried. Types with unknown (italics fonts) or mixed (no italics) expression were placed in both the positive and negative groups (with superscripted numbers indicating the level in the flow chart where their expression was undetermined). Expression data for 10 markers can distinguish most of the neuron types (red outlined boxes); however, three groups could not be distinguished (red boxes): Ivy that are SOM-positive and BP, OA and OBP, and B-CCK (if they are NPY-negative, which is presently unknown) and HB that are PV-negative.

We approached this exercise from two directions. One assessed expression of the markers in the neuron types in the order of their usefulness rating, starting with PV (Figure 3). This approach is analogous to what might be followed in an experimental setting when no marker expression data is known. The second approach took advantage of the known marker information presented in this work. Ultimately, both approaches lead to the same result; the difference is in the number of markers needed to reach the end. With the first approach, twelve markers were needed to resolve all but one branch of the identification tree for CA1 SO neuron types (data not shown). Resolving the CA1 Basket (B) neuron type from CA1 Horizontal Axo-axonic (HAA) requires assessing mu opioid receptor (MOR: second to last branch in Figure 5), the 23^rd^ most useful marker according to our metric (Figure 3).

Using approach two, we examined each divided group of neuron types for the best marker to separate them. We began subdividing by NPY because evidence suggests that six of the 17 neuron types are positive and eight are negative for this marker (Figure 1 and Hippocampome.org). For one of the remaining three neuron types, NPY expression is mixed, and for the last two, expression is unknown. These latter three that do not have unequivocal positive or negative expression are placed in both the NPY-positive and NPY-negative groupings.

SOM was chosen as the second marker, and evidence suggests that only CA1 Ivy in the NPY-positive group has mixed expression. The subgroups resulting from the interrogation of SOM expression could not be neatly divided by one marker in the third level of the flow chart (Figure 5). Taking each group separately, PV, CR, and 5HT-3 demonstrated greatest effectiveness in discriminating the neuron types. Similarly in the final two levels, each group was considered independently, and different markers were used to split them.

The final result suggests that the following ten molecular markers can distinguish most of the 17 known neuron types in CA1 SO: NPY, SOM, PV, CR, 5HT-3, nNOS, CB, CB1, CoupTF II, and MOR. In the end, the CA1 Oriens-Alveus (OA) neuron type could not be distinguished from CA1 Oriens Bistratified Projecting (OBP), nor could CA1 Back-Projection (BP) from CA1 Ivy neurons that are SOM-positive. In addition, CA1 Horizontal Basket neurons that are PV-negative cannot be distinguished from CA1 Basket CCK-positive if CA1 Basket CCK-positive is NPY-negative, which is currently unknown.

### Using molecular marker information to group neuron types

In addition to distinguishing neuron types, biochemical profiles also can be useful in grouping neuron types according to their molecular similarities. To accomplish this, we performed two-step cluster analysis on the molecular marker expression data. We analyzed results obtained from five to 20 clusters using from the top eight most useful markers (Figure 3, PV through NG) to the top 20 (PV through substance P receptor), plus a series of analyses using all of the 97 markers in Hippocampome.org. Then, we evaluated the quality of the 224 clustering results by quantifying the tightness and separation of neuron types groups using the ratio of the average marker profile similarities between and across clusters (Calinski-Harabaz index).

The best clustering was generated when 11 markers were used to produce eight clusters. This configuration produced a ratio of between-cluster/across-cluster similarities of 9.7; for comparison, the worst clustering had a value just above 2.0, which is considered the minimum threshold for acceptable quality). Interestingly, when all 20 top markers were used, the optimal number of cluster remained constant and only two neuron types “jumped” class, producing a between-cluster/across-cluster similarity ratio of 4.2. We ordered the clusters by the product of two averages: the pairwise comparison similarity and the number of markers compared. Figure 6 shows the first seven of the eight clusters, accounting for 74 of the 122 neuron types. The eighth cluster is a catchall for neuron types with very limited marker expression information: the pairwise comparisons for the 48 neuron types in cluster 8 compared on average only two markers.

**Figure 6.**
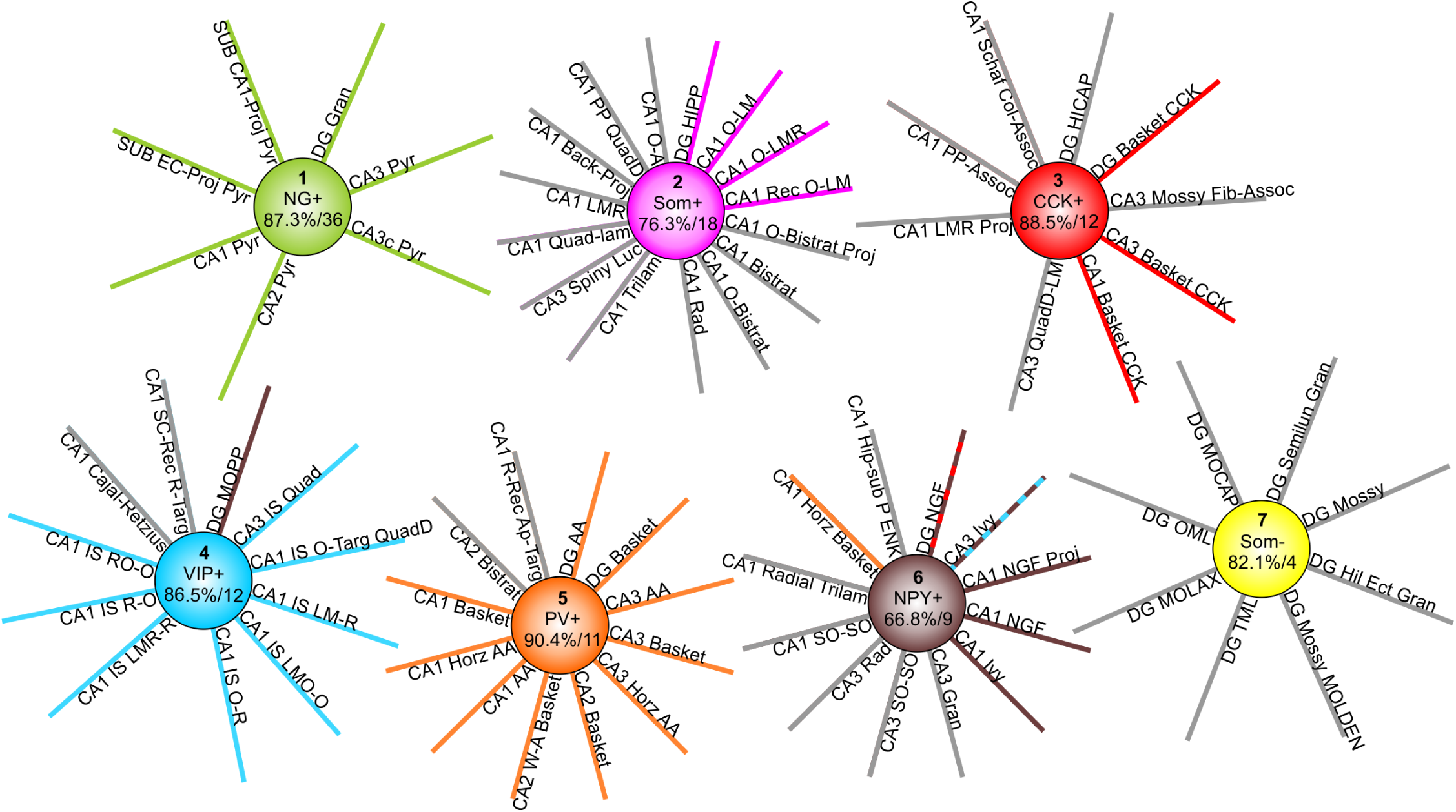
Molecular clustering of Hippocampome.org neuron types. Seven of the eight clusters account for 78 of 122 neuron types. For each cluster, the marker expression interpretation that strongly influences the formation of the cluster is shown in the center along with the average proportion of markers in common and the average number of markers compared across all pairwise comparisons within the cluster. Spokes with the same color as the cluster correspond to “expected” neuron types based on existing biological knowledge. The single spoke of discordant color (e.g., DG MOPP) indicates expected grouping with a different cluster. The two spokes with alternating colors indicate the alternative clustering obtained when using 8 markers instead of 20.

The first cluster contained all and only glutamatergic principal neuron types in DG, CA1, CA2, CA3, and SUB. This group was substantially close molecularly as indicated by an average similarity of 87.3% in pairwise comparisons over an average of 36 markers. A key marker for cluster 1 is the presence of NG, whose expression in the hippocampal formation may be limited to these neuron types (Singec et al., 2004). The second cluster contains most of the O-LM-like neuron types and evidence suggests that all of the neuron types in this group express SOM. A few lesser-known variants of the O-LM morphological family (CA1 OR-LM, CA3 R-LM, and CA3 O-LM), for which SOM presence has yet to be established, are found in cluster 8. Conversely, Cluster 2 contains several SOM-positive types that display different morphological phenotypes, such as CA1 bistratified and trilaminar cells and CA3 spiny lucidum interneurons.

The third category includes CCK-expressing basket cells from all hippocampal subregions in which they have been identified (DG, CA3, and CA1). Cluster 3 also contains CA1 Schaffer Collateral Associated, CA1 Perforant Path Associated, and DG HICAP neuron types, all of which express CCK.

Cluster 4, driven by VIP-positivity, has almost all the interneuron-specific (IS) interneuron types (the single missing cell, CA3 IS oriens, lacks molecular information altogether) as well as DG MOPP cells and CA1 Cajal-Retzius cells, though this latter neuron type, along with CA1 IS O-R, have unknown VIP expression.

The arguably better known types of basket cells are located in the fifth cluster. Every neuron type in the fifth cluster expresses PV, including all known fast-spiking, perisomatic-targeting hippocampal interneurons (classic Basket and Axo-axonic cells) with the sole exception of SUB axo-axonic cells for which no molecular expression information is available.

Cluster 6 is less molecularly homogeneous, with an average similarity of 66.8%. Eight of the 12 neuron types in this cluster are NPY positive, but the others are NPY-negative. CoupTF II is also a driver of this cluster (six of 12 neuron types positive, one negative, one mixed, and four unknown). Notably, this cluster also includes CA1 horizontal basket cells because of their ‘mixed’ expression of both PV and CCK; this is very likely due to the presence of distinct subtypes as in their more common morphological variants of perisomatic-targeting interneurons, which have dendrites in multiple layers.

Lastly, Cluster 7 is driven by the lack of expression of two markers, SOM and mGluR1a. Despite an average similarity of 82.1%, it constitutes a weaker grouping because of the lower average of compared molecular markers (only four). Interestingly, it consists exclusively of dentate cells, including all non-projecting glutamatergic neurons: two types of mossy cells and minor variants of granule cells.

The 8-marker and 20-marker clustering results only differed in the placement of DG Neurogliaform cells (this neuron type has axons that extend into SUB) and CA3 Ivy cells. The 8-marker clustering grouped the former with CCK+ interneurons and the latter with VIP+ cells. Interestingly, adding the information on 12 additional markers was sufficient to shift these two neuron types to the cluster of CA1 Neurogliaform, CA1 Neurogliaform Projecting, and CA1 Ivy cells. Although neither clustering completely matched biologically-intuitive expectations, the lack of sufficient expression data explains most deviations as noted above. The only apparent exception concern DG MOPP cells, which are often associated with the Neurogliaform/Ivy family (Armstrong et al., 2011), as both DG neurogliaform neurons and MOPP neurons have axons and dendrites localized to the outer portion of DG stratum moleculare. In contrast, our results placed DG MOPP in cluster 5, driven by VIP expression (Figure 6).

## Discussion

Hippocampome.org classifies rodent hippocampal neurons based on their primary neurotransmitter and the presence or absence of their axons and dendrites in the respective layers of each sub-region in the hippocampal formation. Upon this foundation is layered information about the proteins or genes each type expresses. Thus, this resource is ideally positioned to help link neuronal morphology and gene/protein expression. An abundance of gene expression data for neurons is on the horizon with the advent of techniques including single-cell RNA sequencing (Poulin et al., 2016; Harborn, Chronister, & McConnell, 2016). Hippocampome.org provides provisional molecular profiles for its neuron types that can be useful to benchmark forthcoming single-neuron transcriptome data from ongoing large-scale studies.

With the goal of maximizing currently available expression data for neurons, we used both neuron type-specific evidence and REIs to formulate interpretations about whether a gene is positive, negative, or exhibits mixed expression in a neuron type. REIs are products of all experiments that analyze expression of multiple genes in a population of cells; consequently, a wealth of them exists in published articles and laboratory records. As this work shows, there are benefits to systematically collecting them. In addition to augmenting the molecular profiles for the neuron types, REIs can bolster our confidence in particular interpretations and highlight weak interpretations based on neuron type-specific evidence, which often comes from very few neurons. Moreover, compiling REIs offers a perspective on co-localizing and mutually exclusive genes. Mechanisms and consequences of these strong expression relationships may point to master transcriptional regulators (Huang, 2014).

REIs hinge on two proportions: the fraction of cells within a neuron type for which the hypothesized relationship would hold; and the probability that the observed relationship is not due to chance. Here, we set the thresholds for use at 66% and 95%, respectively. These values represent a compromise between the number of applicable inferences and the demand that the inferences should apply to a clear majority of neurons within the scope of the REI. Still, with this approach neuron types or subsets of types with small numbers of neurons could be misinterpreted. Data from studies quantifying the number of neurons in each type will help refine these interpretations, not only by identifying types with smaller populations, but also by enabling the analysis of other sources such as *in situ* hybridization data in the Allen Institute Mouse Brain Atlas (Lien et al., 2007).

Our analysis of the molecular expression profiles of the 122 neuron types in Hippocampome.org v.1.3 supports that examining eight or more markers is needed to distinguish the neuron types in the rodent hippocampal formation. Analyzing this number of markers in each cell is likely at or beyond the bounds of traditional techniques such as *in situ* hybridization and immunohistochemistry; consequently, techniques like single cell RT-PCR, RNA sequencing, and RNA and protein arrays may be more useful. The goal of obtaining a molecular identity for all neurons would be greatly assisted by consensus in the neuroscience community on ten molecular markers to be tested in each reconstructed neuron. The most efficient selection process should take into account the amount of information currently known for a marker and the corresponding overall ratio of positive-to negative-expressing neurons. Our analyses suggest that PV, GABA-Aa1, mGluR1a, CR, and SOM are leading candidates. This work, however, is limited to rodent hippocampal neurons; therefore, data from additional brain regions and species are needed. Fortunately, with the advent of high-throughput techniques capable of providing gene expression data in addition to morphology and electrophysiology, such comprehensive information for numerous genes may soon be within reach (Macosko et al., 2015; Fuzik et al., 2016).

Evidence suggests that hippocampal neuron types determined by morphology or electrophysiology overlap substantially (Hosp et al., 2014; Cossart et al., 2006). The extent to which this concordance will extend to molecularly-determined types is unknown, although genome-wide expression data also showed significant overlap (Subkhankulova et al., 2010; Cadwell et al., 2016). Moreover, morphologically similar neurons can have different biochemical content (for example, PV-positive and CCK-positive Baskets in CA1). In Hippocampome.org, we treat molecularly differing neurons with the same axo-dendritic pattern as different types if they differ in their primary neurotransmitter (i.e. glutamate or GABA), or when they display multiple consistent differences in electrophysiology and/or molecular markers. It is possible that the currently most tested molecular markers relate to function rather than morphology, leaving genes responsible for morphological features in the yet-to-be known category. As a result, the molecular marker profiles may better align with the overall function of the neuron type, such as interneuron-specific inhibitor or perisomatic-targeting inhibitor (Kepecs & Fishell, 2014).

A caveat of using molecular profiles to classify neurons is that some genes may be indicative of the state of a neuron rather than a stable marker. Current techniques mark a static moment and typically are lethal to the cells, thus eliminating the opportunity for follow-up data collection. In addition, cells can alter expression of genes in seconds and the amount of RNA and protein present in minutes. Molecularly, neurons even with similar morphological patterns of axons and dendrites may exist on a continuum (Battaglia et al., 2013; Harris et al., 2018). Any markers used to type neurons should be assessed for stable expression.

Another issue that arises when using molecular profiles to type neurons is how many expression differences are necessary to create a new type. Is a single difference sufficient? In the pairwise comparisons presented here, 54 differed by one marker (e.g. CA1 Basket CCK-positive and CA1 Schaffer Collateral Associated, which data to date suggest may differ only by expression of CB out of the 14 markers compared). Morphological characteristics like dendrites in all CA1 layers versus dendrites in two layers and synaptic subcellular targeting preference for the perisomatic region versus dendrites support making these neurons different types; however, without this additional knowledge, what should the decision be?

Finding markers or marker profiles that specifically identify neurons is an important goal. Knowing molecular markers that a particular neuron type expresses will be beneficial in improving our understanding of disorders like epilepsy (Hunt et al., 2013), schizophrenia (Nakazawa et al., 2012; Fung et al., 2010; Brown et al., 2015), and autism (Tatsukawa et al., 2018; Rapanelli et al., 2017), which have been linked to aberrant function of inhibitory neurons. Moreover, it will allow the optogenetic and/or CRE-line targeting of very specific neurons for the purposes of investigation (Bernard, Sorensen, & Lein, 2009; Cardin et al., 2010) and treatment (Hunt et al., 2013; Ghosal, Hare, & Duman, 2017; Chohan & Moore, 2016). Precision is especially important in translational research, and understanding the role of particular neurons in normal and diseased functioning requires a cumulative knowledge incorporating morphology, electrophysiology, and molecular expression profiles (Seung & Sümbül, 2014; Cadwell et al., 2016; Armananzas & Ascoli, 2015; PING, 2008). Hippocampome.org provides a framework to integrate these dimensions and is well positioned to link the forthcoming deluge of hippocampal transcriptome data to morphologically-defined neuron types.

## Author contributions

C.W. and C.R. data mined scientific literature and created computer code for data analysis; C.W., C.R., D.W., and G.A. analyzed data; C.R., D.W., and D.H. engineered and uploaded information to Hippocampome.org; C.W., C.R., D.W., and G.A. wrote the manuscript.

## Acknowledgements

We thank the students who worked in the Facility for Mining the Literature (FaMiLi) and Dr. Alexander Komendantov, a critical member of the team creating Hippocampome.org, for his constructive comments on this work.

This work was supported by the following grants: NINDS (NIH) R01NS39600, NIMH (NIH) U01MH114829, NSF IIS-1302256, AFOSR FA9550-10-1-0385, Keck NAKFI, ONR MURI N00014-10-1-0198.

